# Evaluation of sex and age effects of fiber damage before permeabilization on mitochondrial respiration

**DOI:** 10.1101/2022.11.29.518426

**Authors:** Matthew D. Campbell, David J. Marcinek

**Author notes:** Corresponding author (MDC).

## Abstract

The use of permeabilized muscle fibers (PMF) has emerged as a gold standard for assessing skeletal muscle mitochondrial function. PMF provide an intermediate approach between in vivo strategies and isolated mitochondria that allows the mitochondria to be maintained in close to their native morphology in the myofiber while allowing greater control of substrate and inhibitor concentrations. However, like mitochondrial isolation, the primary drawback to PMF is disruption of the cellular environment during the muscle biopsy and preparation. Despite all the benefits of permeabilized muscle fibers in evaluating mitochondrial respiration and dynamics one of the major drawbacks is increased variability introduced during a muscle biopsy as well as intrinsic variation that exists due to sex and age. This study was designed to evaluate how age, sex, and biopsy preparations affect mitochondrial respiration in extensor digitorum longus, soleus, and gastrocnemius muscle of mice. Here we detail a modified approach to skeletal muscle biopsy of the gastrocnemius muscle of mice focused on maintenance of intact fibers that results in greater overall respiration compared to cut fibers. The improved respiration of intact fibers is sex specific as are some of the changes in mitochondrial respiration with age. This study shows the need for standard practices when measuring mitochondrial respiration in permeabilized muscle and provides a protocol to control for variation introduced during a typical mouse muscle biopsy.

## Introduction

Mitochondria are responsible for meeting the majority of the ATP demand in skeletal muscle (1). As such they play a critical role in skeletal muscle health. Disruption of mitochondrial function occurs in many disease pathologies and also as a natural byproduct of biological aging (2-4). In addition to ATP production mitochondrial oxidative phosphorylation is also a primary source of non-contraction induced oxidant production in skeletal muscle and thus contributes to establishing the redox state in skeletal muscle fibers (5-8) and regulating metabolite and energy homeostasis (9). As skeletal muscle ages the mitochondria tend to become more dysfunctional, disrupting ATP production as well as increasing oxidant production (10, 11).

Multiple approaches are available for assessing mitochondrial respiration that vary in the extent to which they disrupt the interaction between the mitochondria and cell. In skeletal muscle mitochondria are interspersed within the matrix of the myofibrils and into sub sarcolemmal regions (12, 13). *Ex vivo* testing of mitochondrial function is usually performed by isolating mitochondria, however breaking up the myofibrillar matrix leads to disruption of the mitochondrial morphology (14) as well as removal of mitochondria from their biological environment disturbing feedback loops and interactions with the rest of the cell environment (15). In contrast to isolated mitochondria *in vivo* testing of mitochondrial function with spectroscopy approaches maintains the morphology and cellular environment of the mitochondria (16-18). However, the drawback to most *in vivo* testing techniques is the inability to adequately ask and answer mechanistic questions about mitochondrial function. To overcome this, protocols have been developed to use permeabilized fibers in order to maintain mitochondrial morphology by preserving the myofiber lattice while still allowing the ability to stimulate and or stress specific steps or reactions in mitochondria metabolism (19).

The use of permeabilized muscle fibers (PMF) allows the measurement of respiration by titrating in substrates to stimulate specific electron complexes and mechanistic testing by use of specific inhibitors. It is also useful to measure H_2_O_2_ production in parallel with mitochondrial respiration to better evaluate efficiency of flux through the electron transport system. While PMFs have been used for years to evaluate mitochondrial respiration and oxidant production (20, 21) many labs have protocols that deviate slightly from protocols used by other labs and even collaborators. This can contribute to conflicting results and makes replicating studies that much more difficult. Gastrocnemius PMF are the most commonly used skeletal muscle tissue in mice because of the ease of dissection, the heterogeneity of fiber types, and the large amount of tissue generated. However, the gastrocnemius (GAS) presents two obstacles: 1) the GAS is composed of both white (less mitochondrial dense type 2B and X fibers) and red (more mitochondrial dense type 2A fibers) tissue and, 2) the GAS is variably pennate depending largely on the section of muscle used for assays, causing difficulty in isolating intact fibers. One of the most common preparations for mouse GAS fibers is to use cut fibers from the mitochondrial-dense red portion of the GAS (22, 23). In contrast, two other commonly used rodent muscle fibers, the extensor digitorum longus (EDL) and soleus (SOL), are much smaller than the GAS with a different fiber geometry and much more modest angles of pennation (24). Angles of pennation can have far-reaching implications in fiber force development and measurement (Close 1964, Brooks & Faulkner 1988), however for the purposes of fiber biopsies used to measure mitochondria, the angle of pennation seems largely related to the ability to tease fibers apart allowing better permeabilization and increased diffusion of substrates and inhibitors across membranes. Because the EDL and SOL have lower overall pennation compared to the GAS the permeabilized fibers from these muscles tend to be much more intact prior to, during, and following permeabilization. It is unclear how cutting muscle fibers may affect overall function of the mitochondria and thus the ability to detect changes in mitochondrial function between muscle types and with aging and disease. This study tests whether cutting GAS muscle fibers during tissue preparation impacts the ability to detect age related changes in mitochondrial function in male and female mice. Furthermore, we compare the age-related changes observed in cut and intact GAS mitochondria to those of intact EDL and SOL to determine how treatment and preparation of fibers prior to permeabilization impacts comparisons between muscles.

## Materials and methods

### Animals

University of Washington Institutional Animal Care and Use Committee reviewed and approved all experiments performed in this study. We procured male and female C57BL/6 mice from the aged mouse colony at the National Institute of Aging. At the time of experimentation adult male and female mice were 6-7 months old. Aged male and female mice were 26-27 months old. All mice received water and chow ad libitum and were maintained on a 14/10 light/dark cycle at 21°C prior to experimental procedures.

### Tissue preparation, dissection, and measurement of respiration and H_2_O_2_ production

#### Gastrocnemius

Intact GAS was removed and processed as described in results. Following permeabilization one intact and one cut section were flash frozen and saved for follow up measurement of citrate synthase to determine mitochondrial content. Fiber teasing and permeabilization proceeded immediately following dissection.

#### EDL and Soleus

All animals were cervically dislocated and EDL and SOL muscle were removed intact and placed in BIOPS solution (**Table 1**) on ice. Muscles were further dissected parallel to muscle fibers from distal to proximal tendon to produce a bundle of fibers approximately 2-5 mg in mass. Fiber teasing and permeabilization proceeded immediately following dissection. Unused muscle fibers were flash frozen in liquid nitrogen and saved for citrate synthase measurement.

**Table 1.**
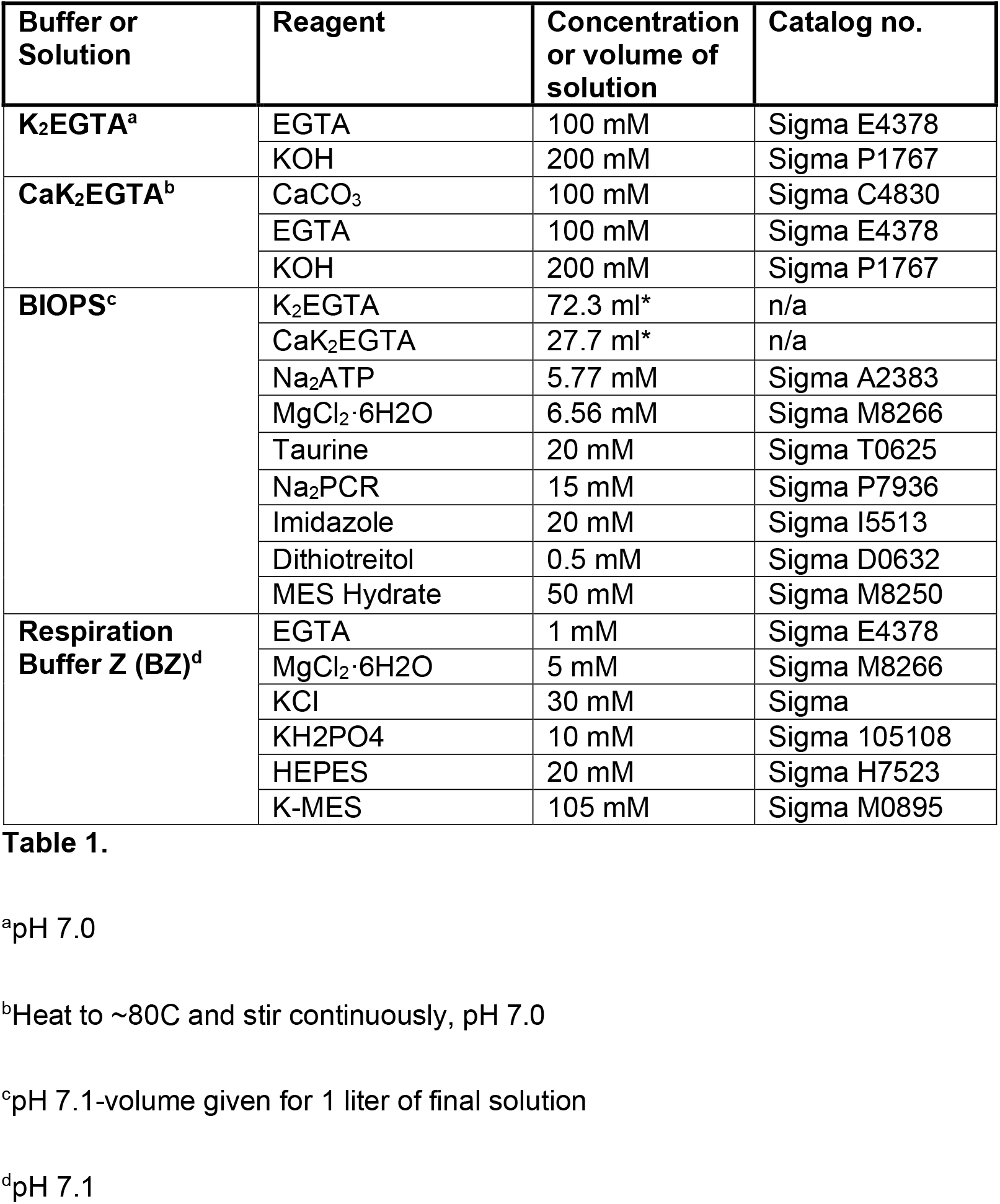
Solution Compositions and Catalog Numbers.

| Buffer or Solution | Reagent | Concentration or volume of solution | Catalog no. |
| --- | --- | --- | --- |
| **K<sub>2</sub>EGTA<sup>a</sup>** | EGTA | 100 mM | Sigma E4378 |
|  | KOH | 200 mM | Sigma P1767 |
| **CaK<sub>2</sub>EGTA<sup>b</sup>** | CaCO<sub>3</sub> | 100 mM | Sigma C4830 |
|  | EGTA | 100 mM | Sigma E4378 |
|  | KOH | 200 mM | Sigma P1767 |
| **BIOPS<sup>c</sup>** | K<sub>2</sub>EGTA | 72.3 ml* | n/a |
|  | CaK<sub>2</sub>EGTA | 27.7 ml* | n/a |
|  | Na<sub>2</sub>ATP | 5.77 mM | Sigma A2383 |
|  | MgCl<sub>2</sub>·6H<sub>2</sub>O | 6.56 mM | Sigma M8266 |
|  | Taurine | 20 mM | Sigma T0625 |
|  | Na<sub>2</sub>PCR | 15 mM | Sigma P7936 |
|  | Imidazole | 20 mM | Sigma I5513 |
|  | Dithiotreitol | 0.5 mM | Sigma D0632 |
|  | MES Hydrate | 50 mM | Sigma M8250 |
| **Respiration Buffer Z (BZ)<sup>d</sup>** | EGTA | 1 mM | Sigma E4378 |
|  | MgCl<sub>2</sub>·6H<sub>2</sub>O | 5 mM | Sigma M8266 |
|  | KCl | 30 mM | Sigma |
|  | KH<sub>2</sub>PO<sub>4</sub> | 10 mM | Sigma 105108 |
|  | HEPES | 20 mM | Sigma H7523 |
|  | K-MES | 105 mM | Sigma M0895 |

<sup>a</sup>pH 7.0
<sup>b</sup>Heat to ~80C and stir continuously, pH 7.0
<sup>c</sup>pH 7.1-volume given for 1 liter of final solution
<sup>d</sup>pH 7.1

### Permeabilization, respirometry, and H_2_O_2_ production

Muscle fiber bundles were manually teased apart and permeabilized in BIOPS solution using saponin (50 ug/ml) at 4°C for 40 min. Following permeabilization muscle fibers were washed in BIOPS solution on ice for 5 minutes, followed by wash in respiration buffer Z (**Table 1**) on ice for 5 minutes, and a final wash in buffer Z on ice for 15 minutes to remove any remaining traces of saponin from permeabilization. Following permeabilization all muscle fibers were blotted dry for 40 seconds using filter paper (Whatman 1442), wet weighed, and placed into a 2 ml chamber of an Oxygraph 2K dual respirometer/fluorometer (Oroboros Instruments, Innsbruck, Austria) at 37°C and stirred gently during substrate and inhibitor titrations.

Respiration was measured as previously described (21). Briefly, substrates, uncoupler, and inhibitors were titrated in the following order and concentrations (see **table 2** for full details): Amplex Ultrared (10 uM), HRP (1 unit/ml), SOD (5 units/ml), H_2_O_2_ (0.1 uM) x2 for fluorescent calibration, malate (2 mM), pyruvate (5 mM), glutamate (10 mM), ADP (2.5 mM), succinate (10 mM), cytochrome C (10 mM), a single titration of FCCP (1.0 uM), rotenone (0.5 uM), antiymycin A (2.5 uM), ascorbate (2 mM), TMPD (0.5 mM), KCN (1 mM). Oxygen signal is allowed to fully stabilize for at least 5 minutes before measurement of oxygen consumption rate. Oxygen concentration was maintained between 200-500 uM. Once oxygen concentration drops below 250 uM chambers are opened and reoxygenated using a small bolus of pure O_2_. Following addition of sample and oxygenation of the chambers, illumination is turned off, amplex red, HRP, and SOD are added to the chambers. Two injections of H_2_O_2_ allowed a three-point fluorescent calibration prior to the start of substrate titrations. Addition of cytochrome C interferes with the amplex red fluorometric signal disallowing oxidant measurements following its addition to the chamber. Runtime for the fluorometric portion of these experiments was usually completed within 30-45 minutes including the time necessary to calibrate the fluorescent signal. Total experimental runtimes typically lasted between 90 and 120 minutes beginning with addition of sample to the chamber and ending with oxygen signal stabilization following addition of KCN. Respiration and oxidant values were also normalized to mitochondrial content measured by citrate synthase activity to evaluate mitochondrial quality vs overall mitochondrial respiration capacity. A representative trace of respirometry and fluorometry experiments can be seen in S1 Fig.

**Table 2.** Reagents, Final Concentrations, and Catalog Numbers.

Substrates, uncouplers, and inhibitors were added in the order presented here
| Reagent | Final Concentration | Catalog | Respiratory State |
| --- | --- | --- | --- |
| Amplex Ultrared | 10 μM | Thermo Fisher A36006 | |
| Horseradish Peroxidase | 1 unit/ml | Sigma P8250 |  |
| Superoxide Dismutase | 5 units/ml | Sigma S8160 |  |
| Malate<sup>a</sup> | 2 mM | Sigma M1000 |  |
| Pyruvate<sup>a</sup> | 5 mM | Sigma P2256 |  |
| Glutamate<sup>a</sup> | 10 mM | Sigma G1626 | State 4/Leak |
| ADP<sup>b</sup> | 2.5 mM | Sigma A5285 | State 3 Complex I |
| Succinate<sup>a</sup> | 10 mM | Sigma S2378 | State 3 Complex I+II |
| Cytochrome C | 10 mM | Sigma C7752 | State 3 MAX |
| FCCP | 1.0 μM | Sigma C2920 | State 3 Uncoupled |
| Rotenone<sup>c</sup> | 0.5 μM | Sigma R8875 | Uncoupled Complex II |
| Antimycin A<sup>c</sup> | 2.5 μM | Sigma A8674 | Non-mitochondrial Rox |
| Ascorbate<sup>d</sup> | 2 mM | Sigma A7631 | Complex IV (+ TMPD) |
| TMPD<sup>e</sup> | 0.5 mM | Sigma T3134 | Complex IV (+ ascorbate) |
| KCN | 1 mM | Sigma 60178 | Complex IV correction |

<sup>a</sup>pH 7.0
<sup>b</sup>Prepare in 300mM MgCl<sub>2</sub>, pH 7.0
<sup>c</sup>Dissolve in 100% EtOH
<sup>d</sup>pH to 6.0 with ascorbic acid
<sup>e</sup>Titrate **after** ascorbate

### Respirometry and fluorometry analysis

Respirometry and fluorometry traces were analyzed using Datlab software version 7.4.0.4 (Oroboros Instruments, Innsbruck, Austria). Following each titration of substrate, uncoupler, or inhibitor the oxygen signal stabilized for ≥ 5 minutes. This stable portion of the oxygen flux trace was marked and the average oxygen consumption per mass was measured. For fluorometry the time points of stable oxygen signal were copied over to the amp flux trace and the average amp flux per mass values were measured that correspond the exact times of stable oxygen flux per mass. Non-mitochondrial residual oxygen consumption (Rox) was measured based on stable oxygen flux levels following addition of antimycin A to inhibit ETS complex III. Reported values of respiratory control states include a correction for this Rox. Additionally, complex IV respiration was calculated by direct stimulation with ascorbate and TMPD followed by inhibition using KCN. The values obtained following inhibition with KCN were subtracted from maximum complex IV flux following addition of complex IV substrates to account for auto-oxidation of TMPD. All oxygen and amperometric flux values were corrected and copied over to Graphpad Prism 9 software for further evaluation and statistical analysis.

### Mitochondrial content

Mitochondrial content was evaluated by measuring citrate synthase activity. Previously snap-frozen muscle fibers (∼5-10 mg) were pulverized using 25 ml ice cold CelLyticMT (Millipore Sigma C3228) containing 0.2% protease inhibitors (Millipore Sigma P8340) and 1% phosphatase inhibitors (Millipore Sigma P2850) in a bullet blender on ice and measured for protein content using a Bradford colorimetric assay (Thermo Fisher 23236). Mitochondrial content was measured using approximately 5-10 mg of tissue with a colorimetric citrate synthase activity kit (Sigma MAK193).

### Statistics

Statistical analysis was performed using Graphpad Prism 9 software. For direct comparisons of cut vs. intact fibers significance was determined using two-tailed students paired t-test as noted in Figure legends. For comparisons of age, sex, or both, significance was determined with one-Way ANOVA including Tukey correction for multiple comparisons, two-way or Mixed-effects repeated measures ANOVA including Sidak corrections for multiple comparisons as noted in Figure legends. Levels of significance and sample sizes are also noted in Figure legends.

### Protocol Link

The protocol described in this peer-reviewed article is published on protocols.io, dx.doi.org/10.17504/protocols.io.bzjap4ie and is included for printing as supporting information file 1 with this article.

## Results and discussion

### Cutting gastrocnemius fibers impacts mitochondrial respiration in male mice

Gastrocnemius muscle is frequently permeabilized and used to measure mitochondrial respiration capacity (22, 23). The myofibers of the mouse GAS are arranged in a complex orientation of varying pennate patterns (25). This arrangement results in variable amounts of damage to the fibers during the typical dissection and may contribute to variation in results from permeabilized muscle fibers (PMF). We set out to test whether the method of dissection affected the mitochondrial function of PMF in young and aged mouse GAS muscles. To establish a more uniform protocol for better repeated measures we took healthy adult and aged male and female mice and dissected out GAS. Immediately following cervical dislocations intact GAS was removed at the proximal and distal tendon and placed in BIOPS (see **Table 1** for composition) solution on ice (Fig 1A). The GAS was further dissected along the midline and approximately 20 mg of intact red GAS spanning a bifurcated tendon was separated for permeabilization (Fig 1B). Remaining white GAS was cut from the opposite side of the tendon leaving only red GAS fibers running approximately perpendicular between opposing tendons (Fig 1C). Red GAS was cut into four sections parallel to the direction of the muscle fibers to keep muscle fibers intact. Individual sections are roughly the same size/weight. For an example of one section see Fig 1D. Following manual teasing and separation of fibers until all fibers were visibly disconnected from parallel adjacent fibers. Two sections were sheared perpendicular to the fibers using surgical scissors along one tendon to produce “cut” fibers, an example of a fiber bundle following teasing but prior to shearing is illustrated in Fig 1E. The other two sections remained attached to the tendon on either side to produce “intact” fibers.

**Fig 1.**
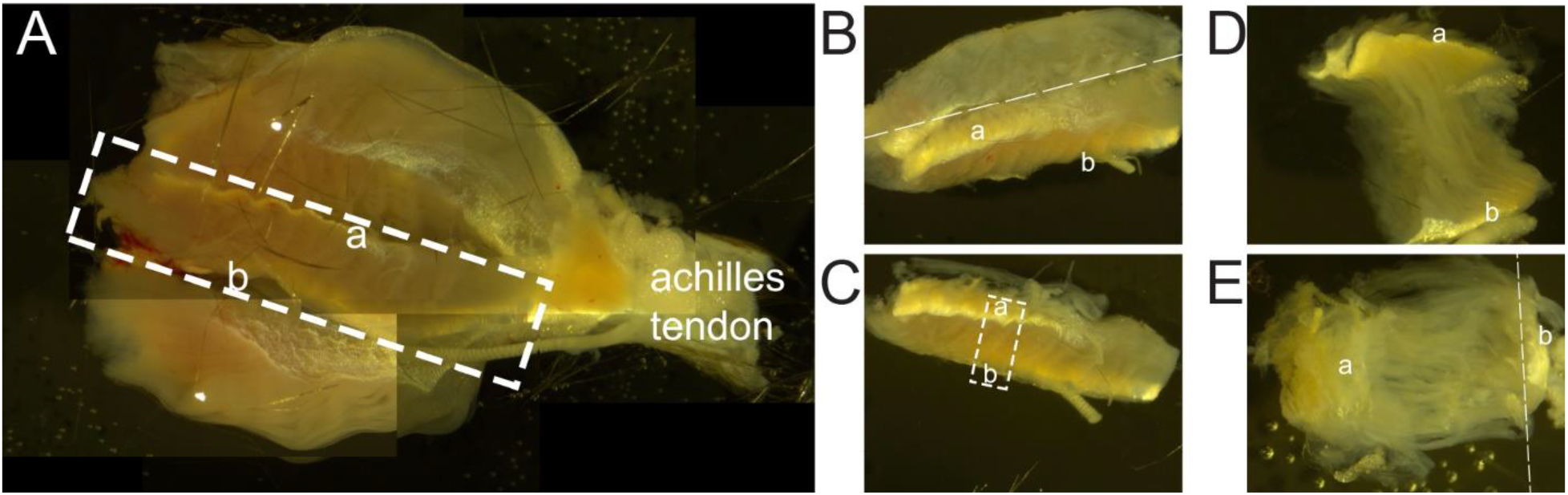
Dissection of mouse gastrocnemius muscle. **(A)** The GAS is removed intact tying ends with 2-0 surgical silk. Reconstructed view is looking down on the internal portion of the GAS normally adjacent to the tibia. Achilles tendon is labeled in white for orientation. White bounding box indicates area of the GAS that is removed keeping tendons labeled **a** and **b** intact. **(B)** Liberated, intact red GAS rotated along the sagittal plane revealing white GAS still attached to the posterior portion of the red GAS opposite tendons **a** and **b**. Cutting proceeds along the dashed white line. **(C)** Red GAS is cut into sections parallel to the direction of muscle fibers running between the tendons labeled **a** and **b. (D)** The final biopsy of red GAS prior to manual teasing showing intact fibers connected to opposing tendons **a** and **b. (E)** Biopsy after gentle teasing of the muscle fibers showing fibers separated but still intact connected to opposing tendons **a** and **b**. Shearing along the indicated dashed white line would produce “cut” fibers.

Permeabilization proceeded as follows: (1) fibers were permeabilized in BIOPS solution containing saponin (50 ug/ml) for 40 minutes on ice, (2) washed in 1 ml BIOPS for 5 minutes on ice, (3) washed in 1 ml respiration buffer Z (BZ, see **table 1** for composition) for 5 minutes on ice, and (4) finally washed for 15 minutes in BZ on ice. Following all wash steps fibers were blotted dry for 40 seconds on filter paper (Whatman 1442) and wet weighed. One intact bundle and one cut bundle were flash-frozen in liquid nitrogen for subsequent citrate synthase and biochemical assays. The other intact bundle and cut bundle were added to 2 ml chambers of an Oroboros O2K dual fluorometer/respirometer (Oroboros Instruments, Innsbruck, Austria) at 37°C and stirred gently during substrate and inhibitor titrations. Following respiration titration protocols data were normalized to citrate synthase activity assessed from the flash-frozen adjacent fiber bundles to account for potential differences in mitochondrial content between the preparations.

Respiration in PMF from cut fibers of adult male mice was significantly decreased compared to PMF from intact fibers under all respiratory states following addition of saturating ADP and succinate (Fig 2A). However, despite changes in mitochondrial respiratory capacity, H_2_O_2_ production was not affected by the dissection protocol (S2 Fig A). In contrast, PMF from adult female mice showed no difference in mitochondrial respiration between intact or cut GAS muscle fibers (Fig 2B). There was also no change in H_2_O_2_ production between cut and intact GAS muscle fibers (S2 Fig B). Just like in adult male mice, maximum respiration of PMF from cut fibers of aged male mice was significantly lower than intact fibers under respiratory conditions following addition of ADP (Fig 3A). Cut fibers from aged male mice showed no change in H_2_O_2_ production in adult male fibers (S2 Fig C). In aged female GAS there was a general trend of decreased respiration overall, and significant differences in some respiratory control states (Fig 3B) but no changes in H_2_O_2_ production (S2 Fig D) that were significantly different between cut and intact fibers. Changes in respiration or H_2_O_2_ production was not due to increased damage to the mitochondria by cutting fibers during preparation as there was no significant difference in the increase of respiration following addition of cytochrome C between cut and intact fibers (Fig 4 A and B). It should be noted though that respiration increase with cytochrome C was higher in both intact and cut GAS fibers compared to EDL and SOL (Fig 4 C and D) suggesting that both preparations lead to more disruption of mitochondrial structure in the GAS than in EDL and SOL.. To control for the potential differences in mitochondrial content due to loss of mitochondria from cut fibers during the tissue processing we subjected intact and cut aged male GAS fibers to a permeabilization protocol and then flash froze tissue to measure citrate synthase (CS) content. There was no difference in mitochondrial content as measured by CS activity following permeabilization between cut and intact fibers in adult mice (Fig 5A). To test whether age made fibers more sensitive to disruption of mitochondrial function by cutting the fibers we directly compared intact fibers of adult animals to intact fibers of aged animals as well as cut fibers in adult animals with cut fibers in aged animals. Although age does depress mitochondrial respiration in PMF in all states except for Complex II and Complex IV driven respiration, there was no significant difference in the effect of age on cut and intact fibers when compared to young cut or young intact fibers respectively when examining both sexes together(Fig 5B), or with each sex examined separately (S3 Fig A and B). It should be noted that addition of pyruvate at high oxygen concentrations impairs complex IV inhibition (26, 27) by cyanide so complex IV values throughout these experiments may not accurately represent respiratory flux through cytochrome C oxidase. We found no age or sex effects on cutting fibers by comparing maximum respiration capacity through complexes I and II following addition of cytochrome C in cut and intact GAS muscle fibers (Fig 5 C and D).

**Fig 2.**
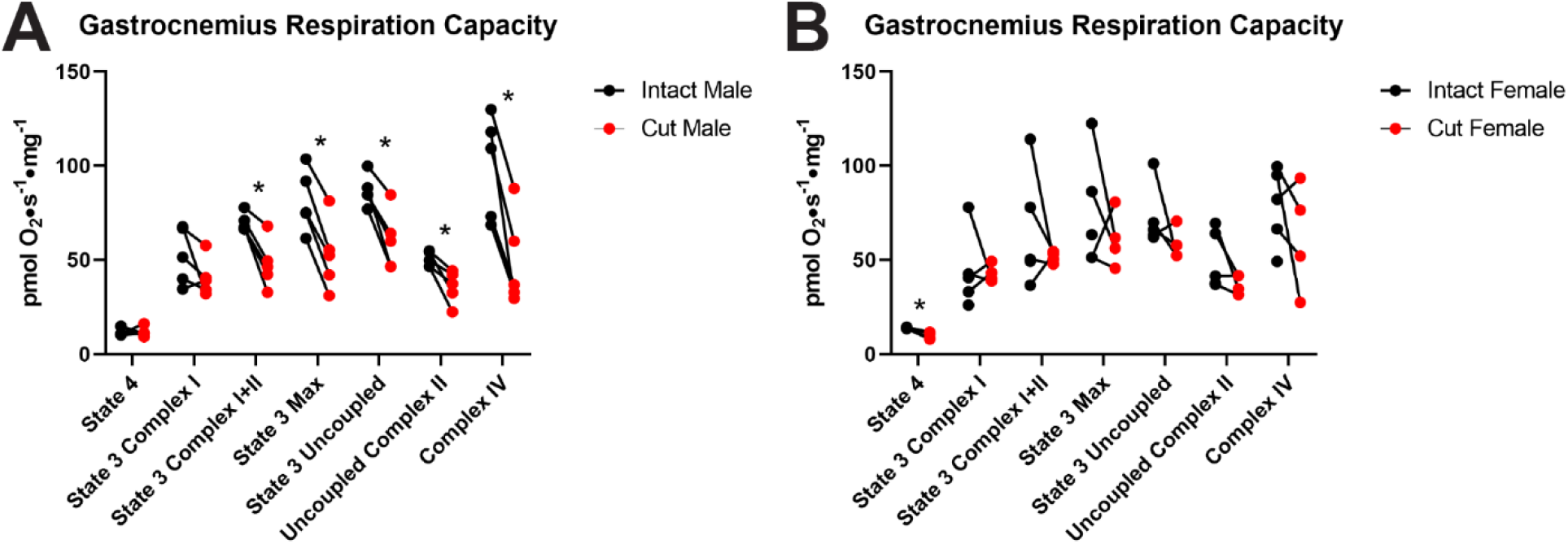
Cut vs intact fibers in adult male and female mice. **(A)** Respiration in adult male GAS fibers. **(B)** Respiration in adult female GAS fibers. All data was analyzed using a paired students t-test. *-significance compared to intact fibers, data represented as mean ± SEM, n=4-5, p<0.05

**Fig 3.**
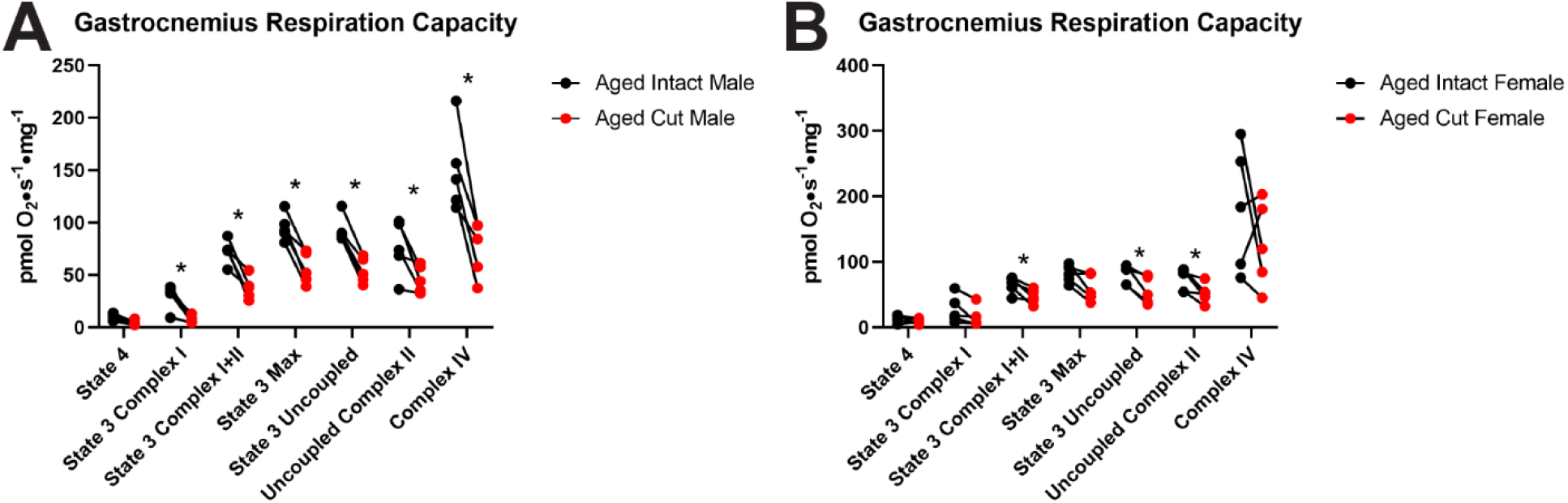
Cut vs. intact fibers in aged female and male mice. **(A)** Respiration in aged male GAS fibers. **(B)** Respiration in aged female GAS fibers. All data was analyzed using a paired students t-test. *-significance compared to intact fibers, data represented as mean ± SEM, n=5, p<0.05

**Fig 4.**
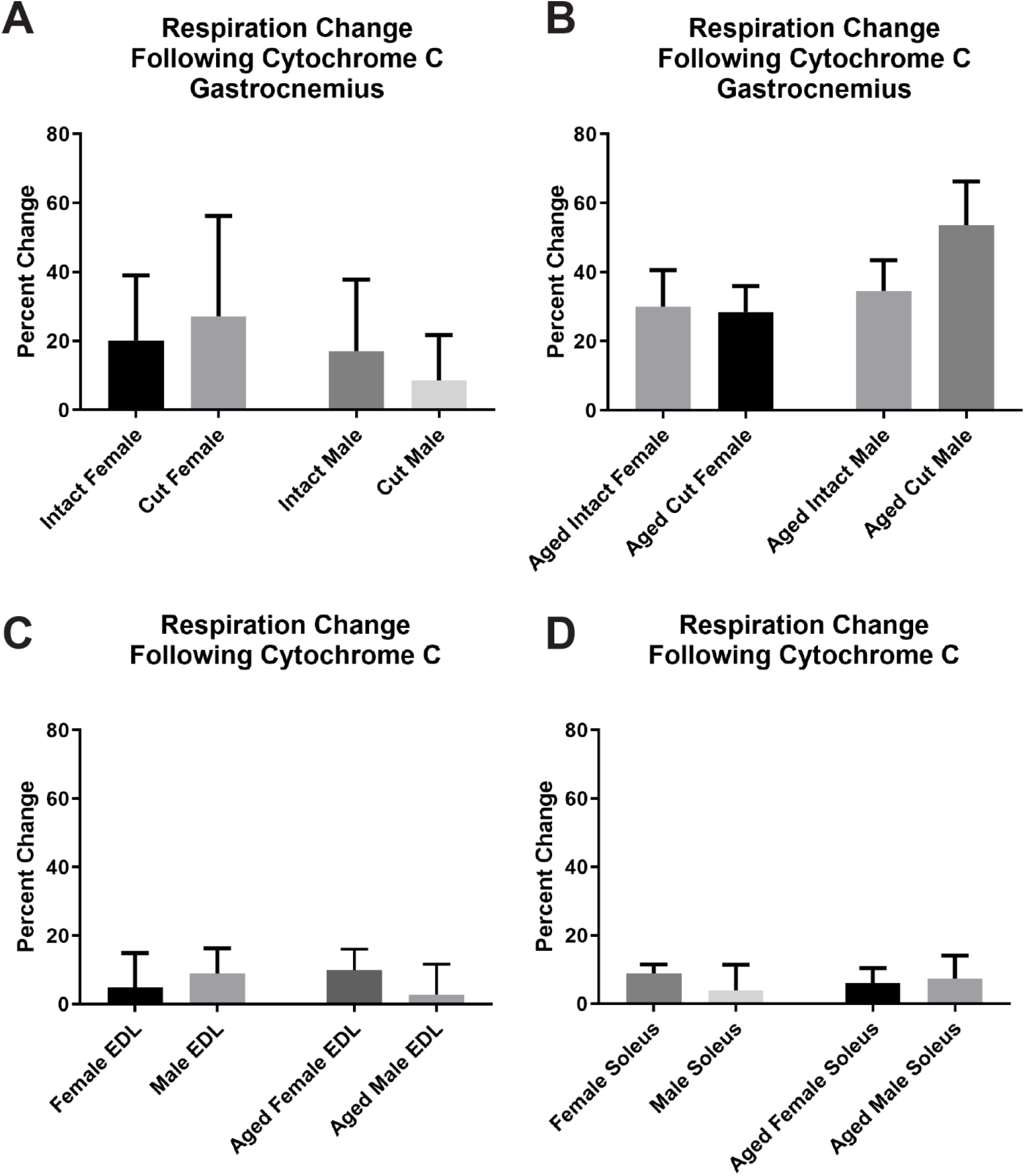
Change in respiration following addition of cytochrome C. **(A)** Adult gastrocnemius **(B)** Aged gastrocnemius **(C)** EDL **(D)** Soleus, all data represented as mean ± SEM, n=5, p<0.05 *significance using two-way ANOVA for all comparisons.

**Fig 5.**
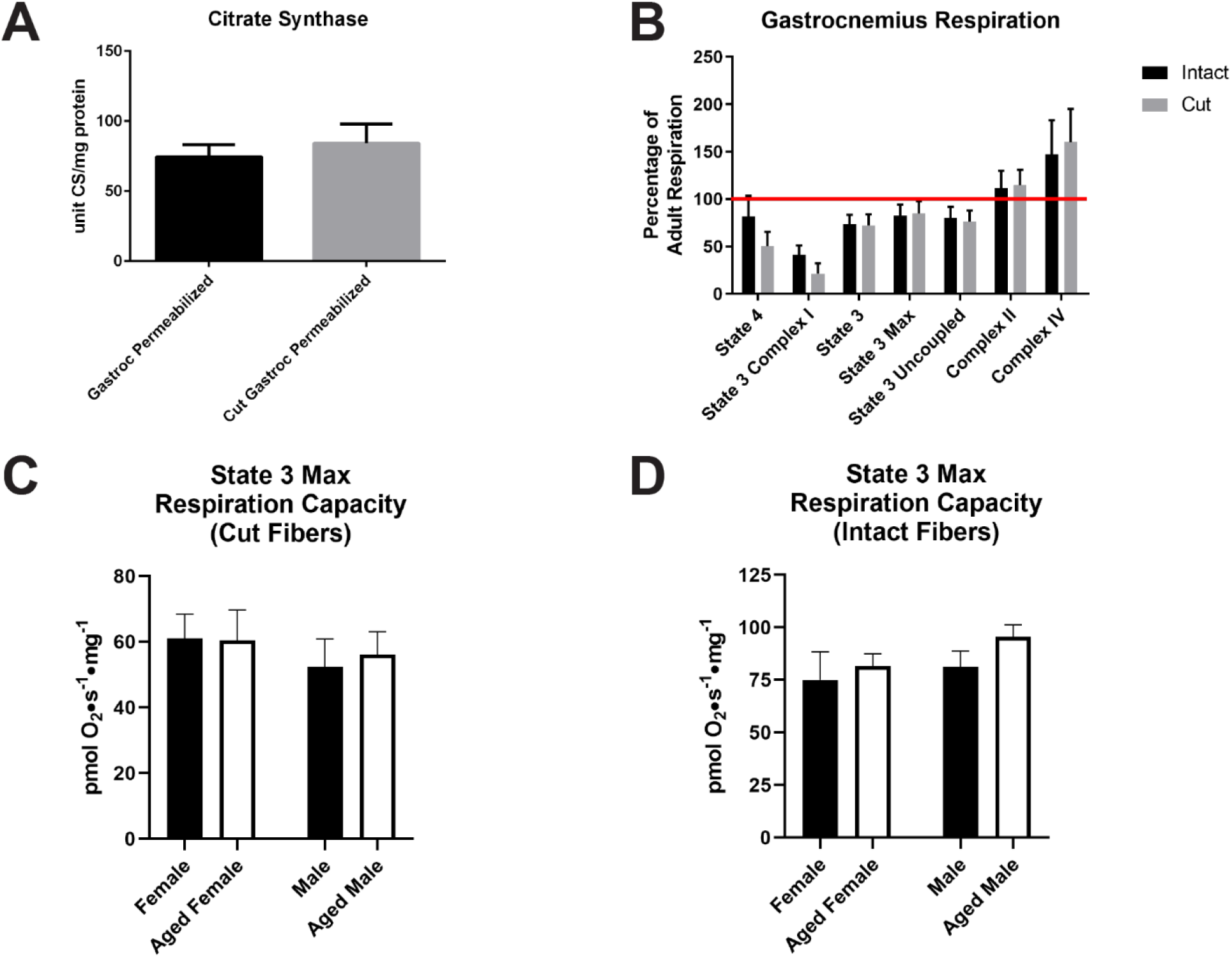
Intact, cut, and sex differences between permeabilized fiber preps. **A**. Citrate synthase activity, n=5 **B**. Respiration of intact vs cut fibers. Respiration in aged intact and aged cut fibers is normalized to adult intact or cut fibers respectively, n=10 **C**. Sex comparison of cut fibers normalized to protein content, n=5 **D**. Sex comparison of intact fibers normalized to protein content, n=5 All data represented as mean ± SEM, n=4-5, p<0.05, *-significance using a paired students t-test for panel **A**, two-way ANOVA for panels **B, C**, and **D**.

### Aging results in decreased mitochondrial quality and increased oxidant production in EDL fibers

The EDL and SOL muscles are frequently used to measure mitochondrial respiration because they have greater proportions of type II and type I fibers respectively, compared to other muscles in the mouse (28). Dissection and permeabilization of EDL and SOL muscles is slightly different from GAS because both muscles can be removed, permeabilized, and bundles of fibers isolated completely intact. EDL muscles were dissected from the legs of healthy adult and aged male and female mice. Fibers in the EDL are arranged with minor pennation to the long axis of the muscle (24) and the entire intact EDL can be dissected from the leg with the fibers intact. Upon excision from the mouse EDL is gently teased apart and intact bundles of approximately 2-5 mg were excised for permeabilization. The remaining unused muscle fibers were flash frozen for follow up biochemical assays. Permeabilization proceeded as follows: (1) fibers were permeabilized in BIOPS solution containing saponin (50 ug/ml) for 40 minutes on ice, (2) washed in 1 ml BIOPS for 5 minutes on ice, (3) washed in 1 ml respiration buffer Z (BZ, see **table 1** for composition) for 5 minutes on ice, and (4) finally washed for 15 minutes in BZ on ice. Following all wash steps fibers were blotted dry for 40 seconds on filter paper (Whatman 1442) and wet weighed before addition to 2 ml chambers of an Oroboros O2K dual fluorometer/respirometer (Oroboros Instruments, Innsbruck, Austria) at 37°C and stirred gently during substrate and inhibitor titrations. Following respiration titration protocols data were normalized to citrate synthase activity to account for potential differences in mitochondrial content between the preparations. PMF were prepared as described in methods and mitochondrial respiration measured in parallel with H_2_O_2_ production. There were no significant differences in maximum respiration capacity of EDL following addition of succinate and saturating ADP to stimulate complex I+II respiration under state 3 conditions (Fig 6A). Following uncoupling and inhibition of complex I by addition of rotenone, aged male and female mice respired no differently than their corresponding adult controls (Fig 6B. H_2_O_2_ production was elevated in EDL of aged male and female mice under state 4, complex I and complex I+II driven state 3 conditions when normalized to mitochondrial content (S4 Fig A and B).

**Fig 6.**
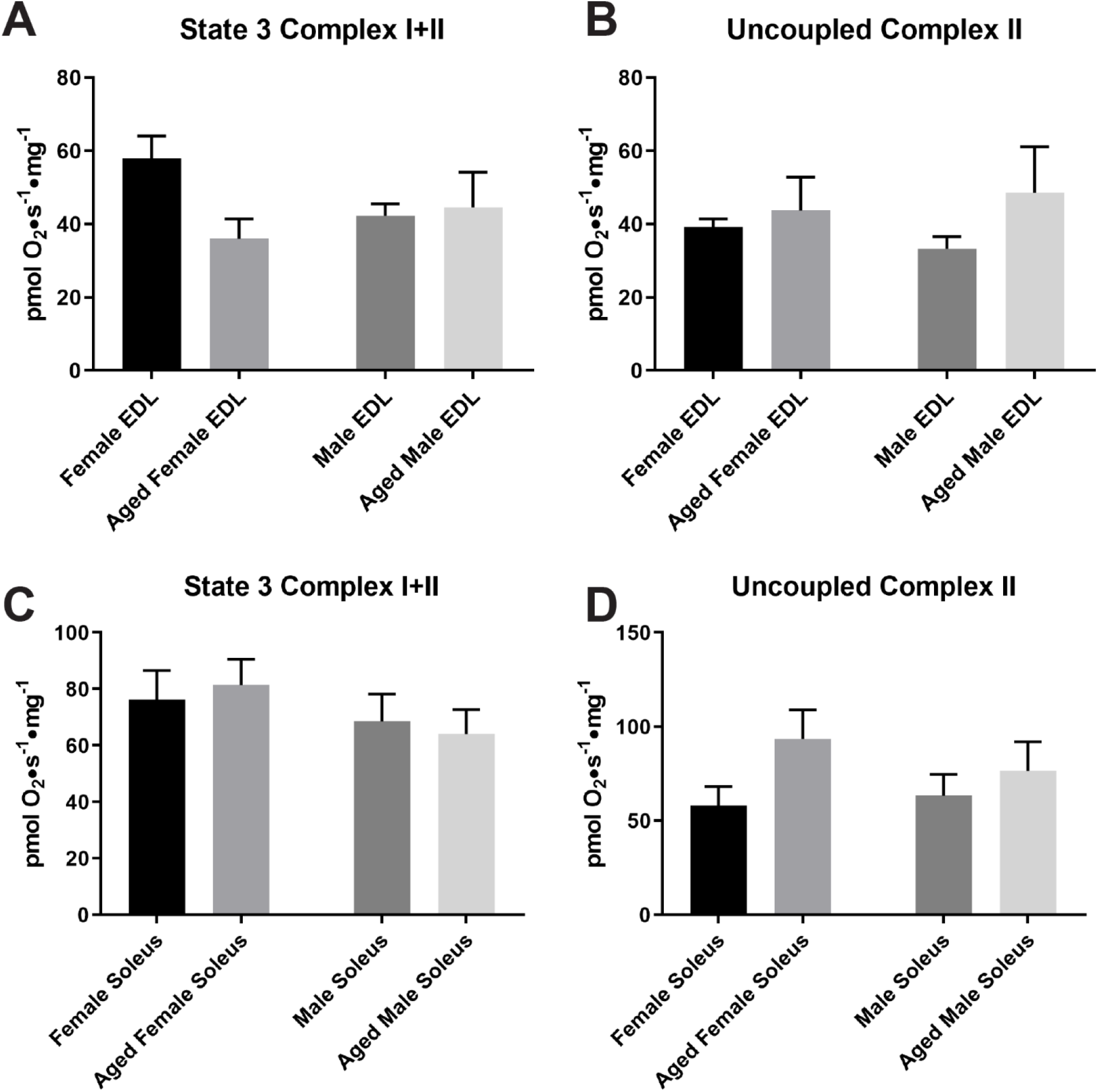
Respiration in the EDL and Soleus of adult and aged female and male mice. **(A)** EDL State 3 respiration after adding succinate and saturating ADP. **(B)** EDL Uncoupled Complex II respiration following complex I inhibition by rotenone. **(C)** Soleus State 3 respiration after adding succinate and saturating ADP. **(D)** Soleus Uncoupled Complex II respiration following complex I inhibition by rotenone. data represented as mean ± SEM, n=5, p<0.05 *significance using two-way ANOVA for all comparisons.

**Fig 7.**
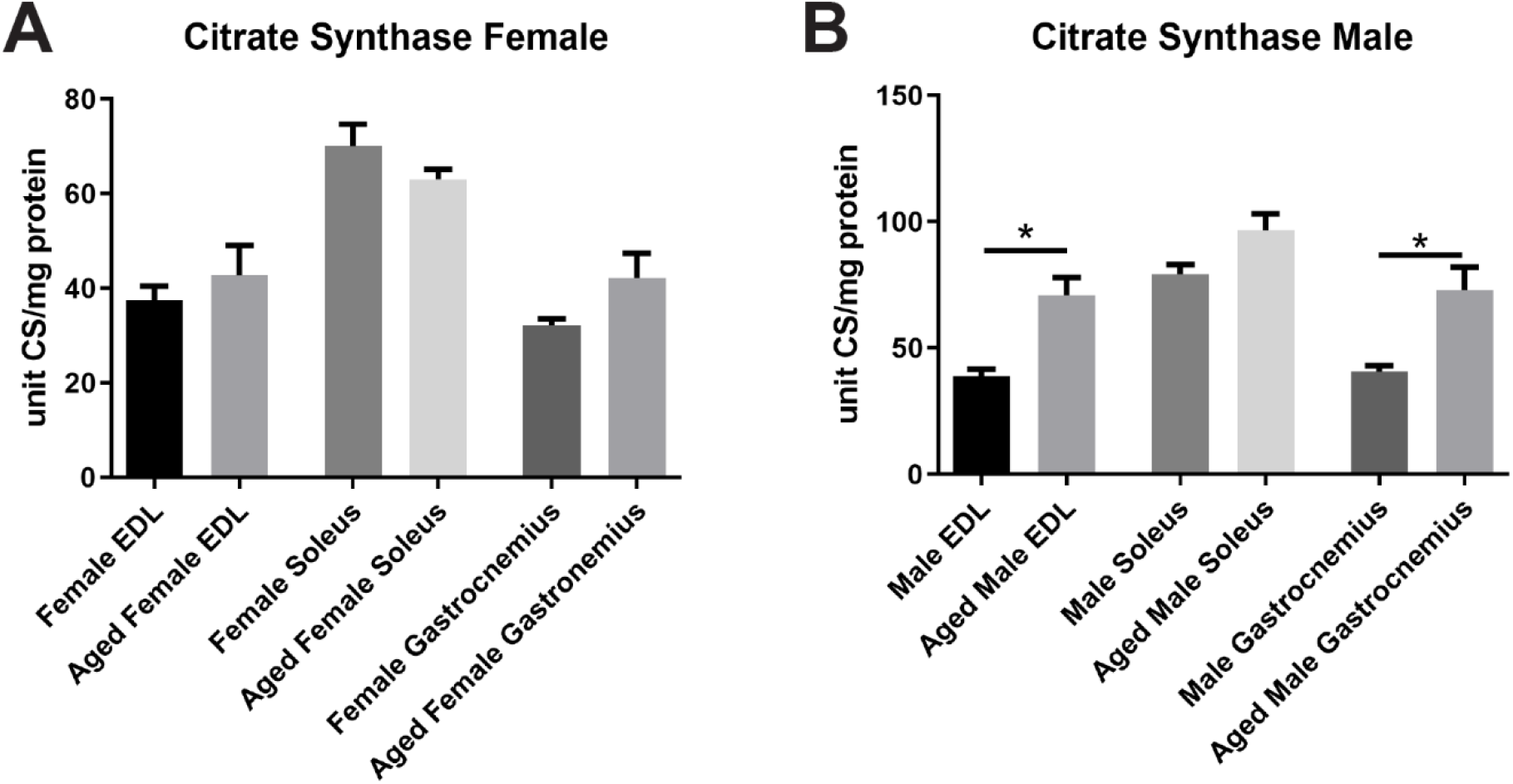
Mitochondrial content of EDL, Soleus, and Gastrocnemius muscle. **(A)** Adult and aged females. **(B)** Adult and aged males. *-significance compared to corresponding adult female or male muscle, data represented as mean ± SEM, n=5, p<0.05 *significance using one-way ANOVA for all comparisons.

### Aging affects mitochondrial respiration and increases oxidant production in Soleus fibers

In contrast to the EDL muscle in mice the SOL muscle is composed of more type I fibers (28). We took healthy adult and aged male and female mice and dissected out SOL muscle. As in the EDL, fibers in the SOL are arranged mostly parallel to the long axis of the muscle and the intact SOL can be dissected from the leg with fibers completely intact. Muscle fibers were treated as described in methods to measure mitochondrial respiration in parallel with H_2_O_2_ production in PMF. Upon stimulation with succinate and saturating ADP neither sex demonstrated a change in respiration (Fig 6C). Uncoupled complex II respiration was also unaffected by age (Fig 6D).Aged males show no significant difference in H_2_O_2_ production compared to adult male controls, however aged female mice do have a significant increase in H_2_O_2_ production compared to adult female controls during state 4, complex I, and complex I+II driven state 3 respiration, while (S4 Fig C and D).

### Aging affects skeletal muscle mitochondrial content in male mice

There are conflicting reports of whether mitochondrial respiration increases or decreases with age (29-31). This makes it necessary to normalize both respiration and oxidant production to mitochondrial content as the number of mitochondria and quality of mitochondria will determine overall respiration efficiency. We measured citrate synthase activity as an indicator of mitochondrial content in EDL, SOL, and GAS. There was no effect of age on citrate synthase activity in female mice for any of the muscles (Fig 8A). However, in aged male mice citrate synthase activity was elevated in EDL and GAS. (Fig 8B).

## Conclusions

The primary goals of this study were to assess the effect of differing permeabilization preparations on GAS muscle fibers and to compare the effects of age on mitochondrial function in PMF between muscles and sexes. Because the fiber geometry in GAS typically leads to cut fibers and isolation of EDL and SOL is done on intact fibers it was important to test the effects of cutting fibers while also including comparisons between muscle types. It is possible that some of the conflicting results in the literature and even between studies is due at least in part to the protocol used for dissection and permeabilization procedures, particularly in the mouse GAS. This may be due to the pennate organization and overall size of the GAS making isolation and permeabilization of intact fibers more mechanically difficult and more complicated for reproduction than the EDL and SOL. Therefore, it is prudent to identify the most uniform protocol for dissection, permeabilization, and measurement as possible for proper reproduction of these assays and so that measurements of mitochondrial respiration in GAS may be more accurately compared to other muscles like the EDL and SOL. We sought to design a procedure for GAS fiber isolations that would keep fibers intact in a similar manner to EDL and SOL preparations. While the primary steps of this procedure follow other published protocols (21), by adding dissection steps fastidiously noting the direction of muscle fibers and the connections to tendon insertion it is possible to remove mostly bundles of fibers that are intact from one end to another. The use of cytochrome C as a test for mitochondrial outer membrane stability (32) suggests that even disruption of the muscle structure necessary to remove samples for preparations of permeabilized intact bundles of GAS contributes significantly to respiratory defects when compared to muscles such as the ELD and SOL which may be removed completely intact.

The manner in which the fibers are dissected and prepared does have a significant influence on the measurements of mitochondrial respiration in GAS fibers, although these results appear to be sex-dependent and independent of age. In both adult and aged mice GAS muscle fibers from male mice are most affected by fiber cutting. A potential explanation for the lack of effect of preparation technique in female mice is greater resistance to oxidative damage in females. In female rats it has been shown that untrained female mitochondria have a greater resistance to oxidant exposure and lower overall lipid peroxidation and antioxidant enzymes than males (33). Previous studies have shown that increases in female longevity are correlated with decreased oxidant production and overall increased antioxidant defense largely due to the increased levels of estrogen (34). This suggests that similar levels of oxidant production as seen in this study between aged male and aged female fibers will disproportionately affect mitochondrial function in male mice. Denervation models used to induce elevated oxidant production that corresponds to an aging sarcopenic phenotype have shown that the bulk of measured oxidants produced in muscle originate from lipid hydroperoxides and not necessarily from sources in the electron transport chain (35). It should be noted that permeabilized muscle fibers require hyperoxygenation to account for greater oxygen dependence due to diffusion and spatial limitations than that of tissue homogenates or cells (36-38). This hyperoxygenation can produce artificially high levels of reactive oxygen species. Although all samples were prepared and measured under the same conditions this does not eliminate the possibility that cut or intact fibers vary in their response to hyperoxygenation. The assays here also do not differentiate between the sources of oxidant production and lipid hydroperoxides are likely a significant contributor to the H_2_O_2_ signal measured in these experiments. However, cutting fibers only revealed differences in oxidant production in aged male mice. The decrease in oxidant production suggests that H_2_O_2_ production is not a significant contributor to the effect cutting fibers has on mitochondrial respiration. Similar effects of age on respiration in both preparations indicates that age is not a contributing factor to the discrepancy found when fibers are cut. The consistent reduction in respiration in cut fibers from male GAS indicates that keeping fibers intact throughout the dissection process, similar to the way EDL and SOL dissection and permeabilizations are performed, is a strategy to maintain mitochondrial quality and reduce artifact in the analysis of PMF. To reduce variability, it appears to be more effective to use intact fibers similar to preparations of the EDL and SOL. Although cutting fibers does not reduce mitochondrial content compared to intact fibers it should be noted that we are unable to adequately retrieve and process tissue following respirometry/fluorometry so it may be possible that during substrate, inhibitor, and uncoupler titrations mitochondria in cut fibers die or are destroyed at a greater rate than intact fibers.

The EDL and SOL are two muscles of particular interest because within mice these muscles contain the largest proportion of type I fibers in the SOL (∼40-50%) or type II fibers in the EDL (∼80-90%) (28, 39). Mechanical changes in SOL and EDL muscle of mice with age have been known for quite some time (40). Previous work in rats has shown that mitochondrial changes with age are fiber-type specific but do not explain differential atrophy in GAS, EDL, and SOL muscles (41). Despite being two heavily studied muscles for their unique properties there have been relatively few studies that directly examine the difference of mitochondrial respiration between EDL and SOL (42-44) and even fewer that have compared how age (42) or sex affects these changes in mice, although work in rats has shown decreases in mitochondrial respiration per unit with age in GAS, EDL, and SOL muscles (41, 45). Sex differences do appear when looking at the aged response in both EDL and SOL muscles when normalized to citrate synthase, indicating that differences in mitochondrial content with age likely underlie some of the changes in respiration.. The difference in response based on sex may also be linked to the inability of males to respond to oxidative insult to the same capacity as females (33). Citrate synthase data shows that aged males have significantly increased mitochondrial content in the EDL compared to adult males consistent with the idea that the elevated content may be a response to decreased mitochondrial quality, which appears to be greater in male than female mice. These data suggest that the decline in mitochondrial respiration in fast twitch fibers with age is less severe in females than it is in males.

In conclusion, this study demonstrates that disruption of mouse skeletal muscle fibers due to cutting has a negative impact on mitochondrial respiration in PMF. The effects of fiber disruption are independent of age, but sex-dependent in mice. Therefore, it is critical to standardize protocols that minimize disturbance of the muscle fibers during permeabilization in order to improve repeatability and to mitigate confounding factors when measuring mitochondrial respiration and oxidant production across muscles, ages, and sexes.

## Author Contributions

M.D.C, D.J.M. designed experiments; M.D.C conducted experiments and collected data; M.D.C, D.J.M analyzed and interpreted data; and M.D.C and D.J.M wrote the manuscript.

## Supporting information

**S1 Fig.**
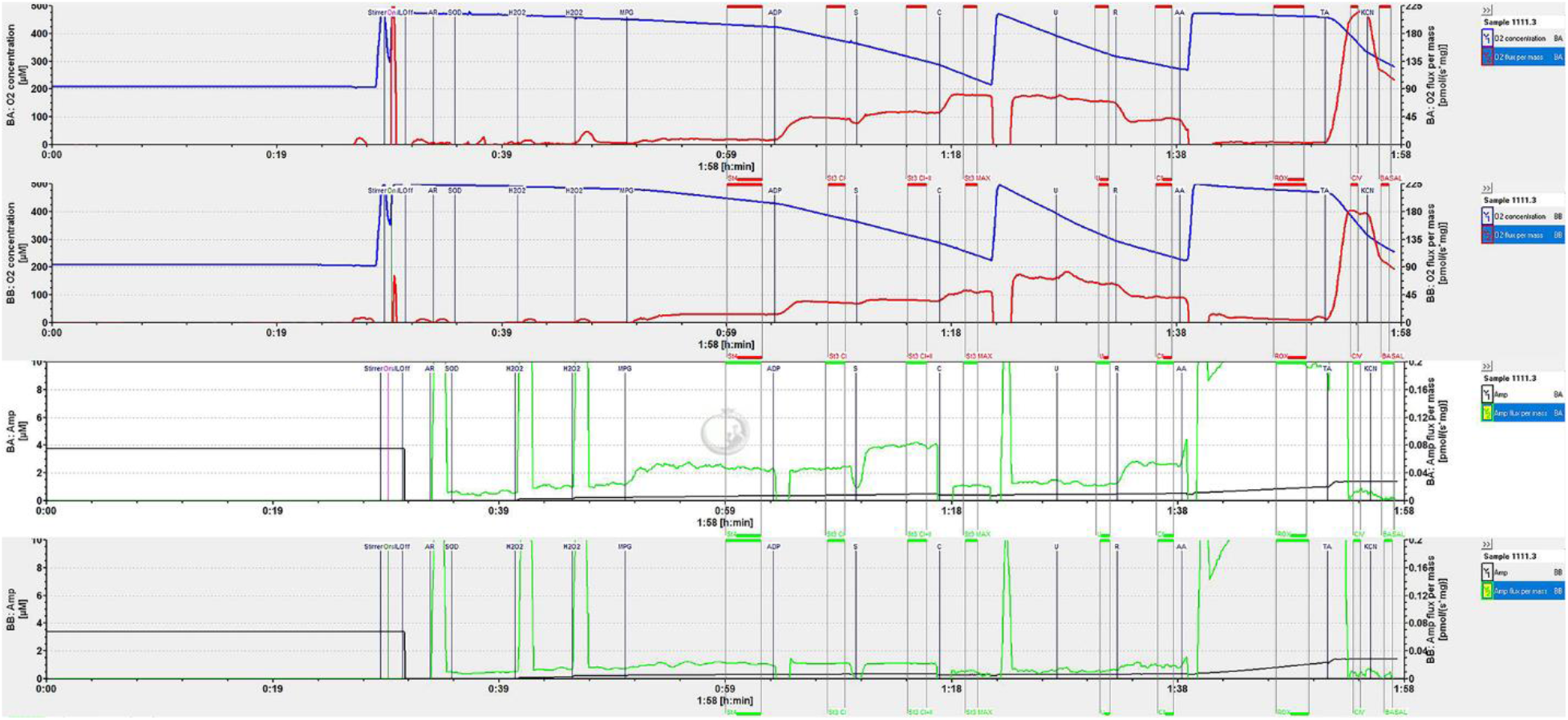
Representative Respirometry and Fluorometry Trace.

**S2 Fig.**
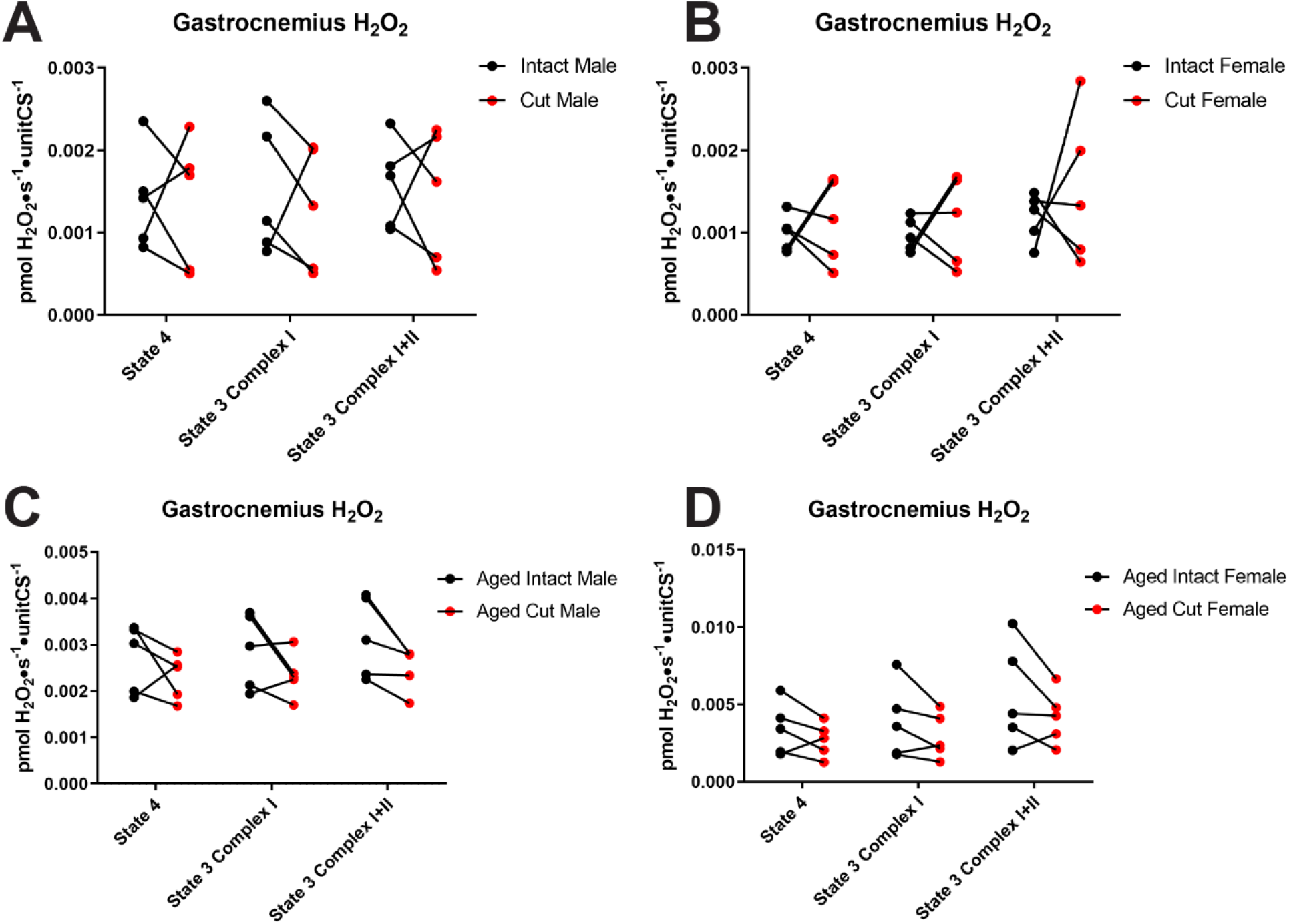
Oxidant production in cut vs intact fibers of gastrocnemius. **(A)** Adult male **(B)** Adult female **(C)** Aged male **(D)** Aged female. All data was analyzed using a paired students t-test. *-significance compared to intact fibers p<0.05, n=5

**S3 Fig.**
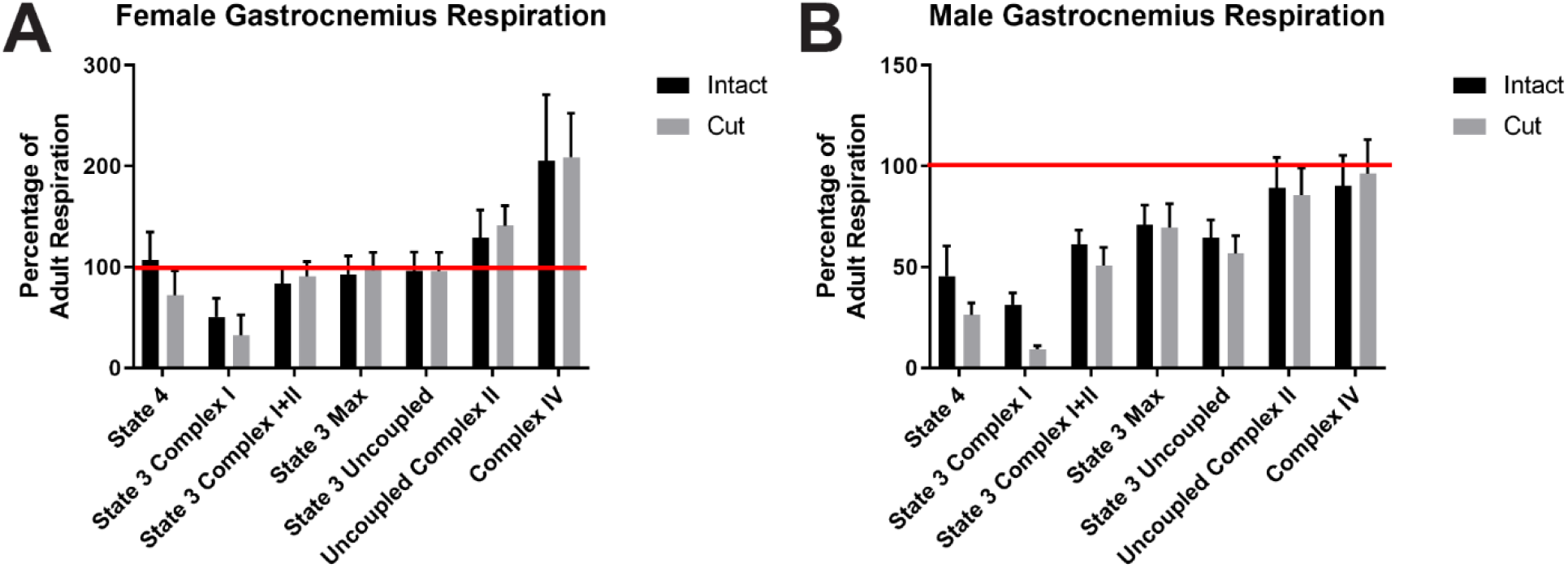
Age effects on cut and intact fiber preps. **(A)** Female intact and cut fibers **(B)** Male intact and cut fibers. All data was analyzed by two-way ANOVA and a Sidak correction for multiple comparisons, n=5.

**S4 Fig.**
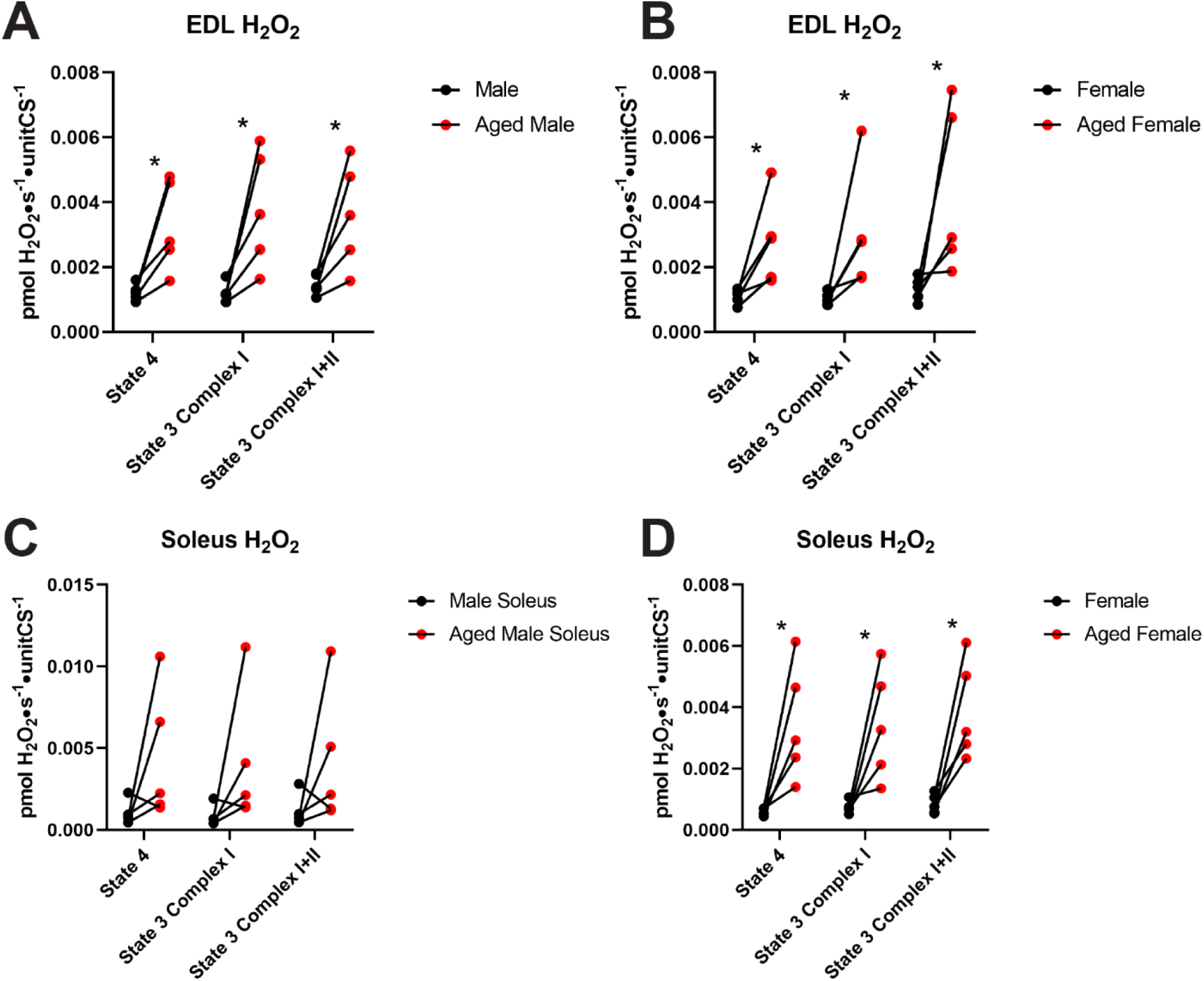
EDL and Soleus oxidant production normalized to mitochondrial content. **(A)** Male EDL oxidant production **(B)** Female EDL oxidant production. **(C)** Male Soleus oxidant production **(D)** Female Soleus oxidant production. All panels were analyzed using students t-test. Panels p<0.05, n=5 *-significance compared to adult controls.

